# DNA methylation-based sex classifier to predict sex and identify sex chromosome aneuploidy

**DOI:** 10.1101/2020.10.19.345090

**Authors:** Yucheng Wang, Eilis Hannon, Olivia A Grant, Tyler J Gorrie-Stone, Meena Kumari, Jonathan Mill, Xiaojun Zhai, Klaus D McDonald-Maier, Leonard C Schalkwyk

## Abstract

Sex is an important covariate of epigenome-wide association studies due to its strong influence on DNA methylation patterns across numerous genomic positions. Nevertheless, many samples on the Gene Expression Omnibus (GEO) frequently lack a sex annotation or are incorrectly labelled. Considering the influence that sex imposes on DNA methylation patterns, it is necessary to ensure that methods for filtering poor samples and checking of sex assignment are accurate and widely applicable. In this paper, we presented a novel method to predict sex using only DNA methylation density signals, which can be readily applied to almost all DNA methylation datasets of different formats (raw IDATs or text files with only density signals) uploaded to GEO. We identified 4345 significantly (*p* < 0.01) sex-associated CpG sites present on both 450K and EPIC arrays, and constructed a sex classifier based on the two first components of PCAs from the two sex chromosomes. The proposed method is constructed using whole blood samples and exhibits good performance across a wide range of tissues. We further demonstrated that our method can be used to identify samples with sex chromosome aneuploidy, this function is validated by five Turner syndrome cases and one Klinefelter syndrome case. The proposed method has been integrated into the *wateRmelon* Bioconductor package.

## Background

DNA methylation is one of the most-studied epigenetic modifications, which typically occurs in the context of a cytosine-guanine dinucleotide motif (CpG) [1]. DNA methylation plays important roles in the stability and regulation of gene expression in the development and maintenance of cellular identity [2]. The dynamic process of DNA methylation and the plasticity of the DNA methylation landscape make genes responsive to the changes of environmental conditions. Several health and lifestyle factors have been found to be associated with DNA methylation signatures, including childhood disease, tobacco smoke, drug use and poor nutrition [3, 4, 5].

Genome-wide analysis of DNA methylation has now become popular and is growing rapidly, owing to array-based profiling technologies. The two most widely used microarray platforms, Infinium HumanMethylation450 BeadChip (450K) [6] and Infinium MethylationEPIC BeadChip (EPIC) [7], offer broad coverage and precise quantification of DNA methylation levels at roughly 480,000 and 860,000 CpG sites respectively.

Epigenome-wide Association Studies (EWAS) are a powerful way to study the relationships between epigenetic variation and human diseases [8]. Apart from sex chromosomes, thousands of CpG sites on autosomes also show very different DNA methylation patterns between males and females [9, 10]. As a result of this, sex has been considered an important co-variate, when undertaking methylation and phenotype association studies.

Many researchers have submitted their methylation microarray datasets to the Gene Expression Omnibus (GEO). Currently, there are over 100,000 HM450k samples and over 18,000 EPIC samples which are publicly available. Most of these have phenotype annotations accompanying them, thus they can be used by other researchers to perform meta-analyses or as independent references to validate their hypothesis. However, many mismatches have been found between annotations and samples, Toker et al. discovered widespread mislabelling in transcriptomics datasets of GEO [11], Heiss et al. found 25% of the datasets they studied contained sex-mismatched samples, particularly in three datasets, more than 30% of the samples were identified as being mislabelled [12]. A large portion of these discrepancies may stem from data entry errors. Researchers should deal with these sex-mismatched samples carefully; the safest way is to remove them directly before downstream analysis.

Currently, there are several methods which can be used to predict the sex of samples from DNA methylation data. The ‘getSex’ function of *minfi* package estimate sex based on the median values of measurements on the X and Y chromosomes respectively [13]; the ‘estimateSex’ method of *sEst* package estimate sex based on the percentages of beta-values and *p*-value in different intervals [14]; The ‘check sex’ method within the *ewastools* package predict sex based on normalized average signal intensity values on the autosomes and the sex chromosomes [12].

In this paper, we propose a novel method to predict the sex of samples using solely DNA methylation levels. We identify a set of significant sex-associated CpG sites, and perform principal component analysis (PCA) on these sites to obtain a sex classifier, and evaluate our method’s performance across a wide range of human tissues. The proposed sex classifier allows users to attribute sex to un-annotated samples on public databases, and also identify samples with sex aneuploidy.

## Results

### Identifying sex-associated CpG loci

To make our method compatible with both 450K and EPIC, we only included 453,152 probes that are present on both arrays. A two-sample *T* -test was used to identify differentially methylated CpG sites between sexes, after Bonferroni multiple comparison correction, those with *p*-value less than 0.01 were selected as the most significant sex associated CpG sites. As a result of this, we obtain 4345 significantly sex-associated sites. In this study we have chosen a relatively strict threshold, as we aim to capture those most robust features which methylate differently and consistently between the two sex groups across various datasets. As expected, most of the sex-associated sites belong to sex chromosomes, with the majority (4047, 93%)located on the X chromosome (ChrX), and with a total of 284 (6.5%) CpG sites located on the Y chromosome (ChrY). As reported previously [9], some regions of the autosomes also show highly significant sex association, here we also identify 14 sites dispersed throughout the autosomes which significantly associated with sex. As shown in Figure 1a, these sex-associated CpG sites on ChrX are distributed across almost the whole chromosome, and with most of them (3781, 93.4%) associated with higher methylation levels in females compared to males, this is mainly because one X chromosome of the female is inactivated and highly methylated. However, we also observed a small portion of CpG sites (266, 6.6%) on ChrX that have higher methylation levels in males compared to females, one possible explanation is that some genes on the X chromosome can escape X chromosome inactivation (XCI) [15].

**Figure 1.**
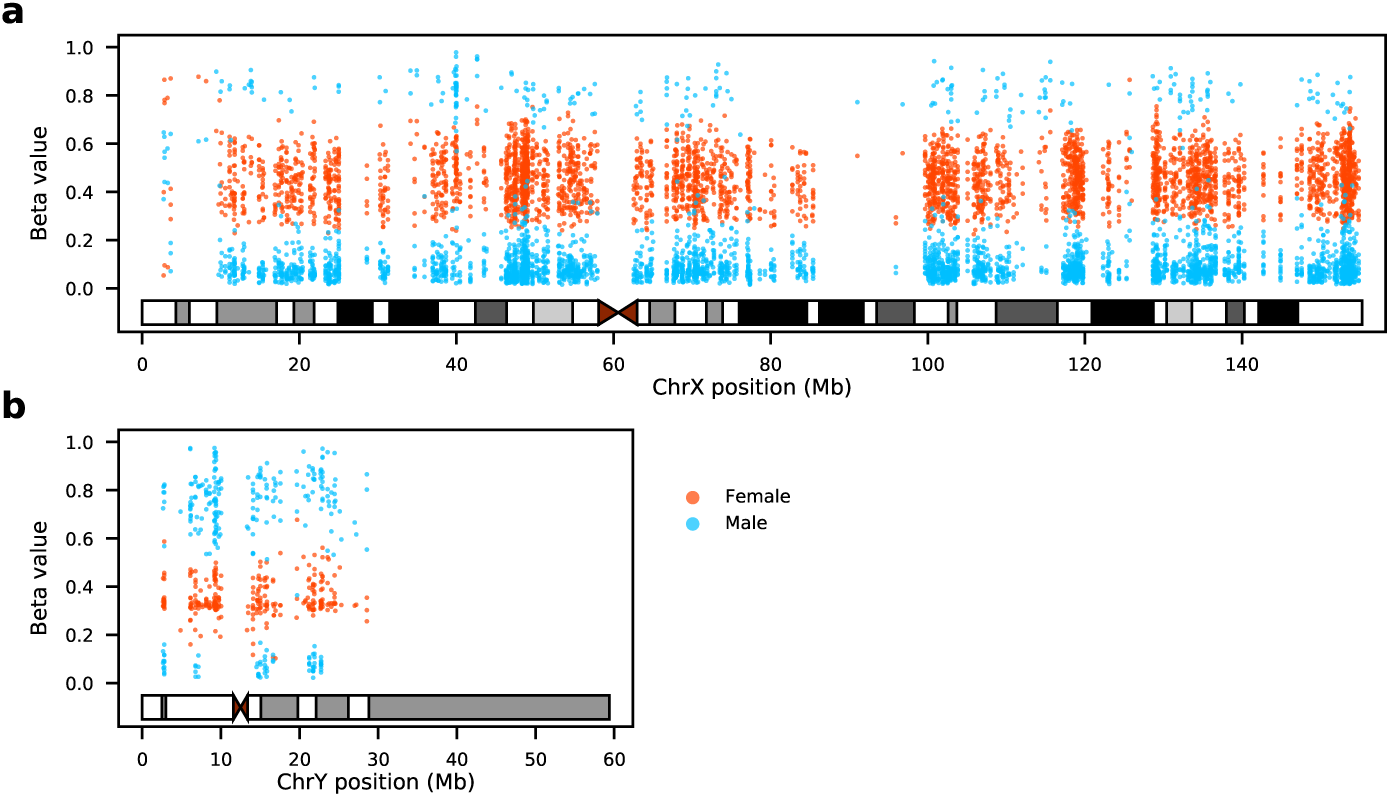
Females and males exhibit distinct methylation patterns at sex-associated CpG sites on the two sex chromosomes. **a**: The X chromosome: most sex-associated CpG sites from females have beta values range between 0.2 and 0.8; most of these sites from males are less methylated (beta values less than 0.2). **b**: The Y chromosome: the identified sex-associated CpG sites of males are highly methylated with beta values greater than 0.6 whereas females exhibited low methylation signals.

Among the 284 sex-associated CpG sites on ChrY, 211 CpG sites have higher methylation levels in male samples (Fig. 1b). Females do not carry Y chromosomes, thus most of the intensity signals of ChrY we observed from females are mainly due to background noise, and the remainders may come from cross hybridisation with similar genomic regions on the autosomes.

### Sex classifier based on sex-associated CpG sites

Since we have obtained a large group of CpG sites which show a significant difference (*p* < 0.01) in methylation levels between males and females, we are able to construct a sex classifier. To begin with, the DNA methylation values of the 4047 sex-associated CpG sites on ChrX from the same training samples are processed using PCA. PCA takes a linear approach to generate reduced dimensions by maximizing the captured residual variance in each further dimension[16]. As shown in Fig. 2a, the first principal component, which explained 98% of total variance, has captured the most sex differences among the all trainging samples. Thus, we could use this first component to separate samples into two categories: 1) with two X chromosomes and 2) with a single X chromosome.

**Figure 2.**
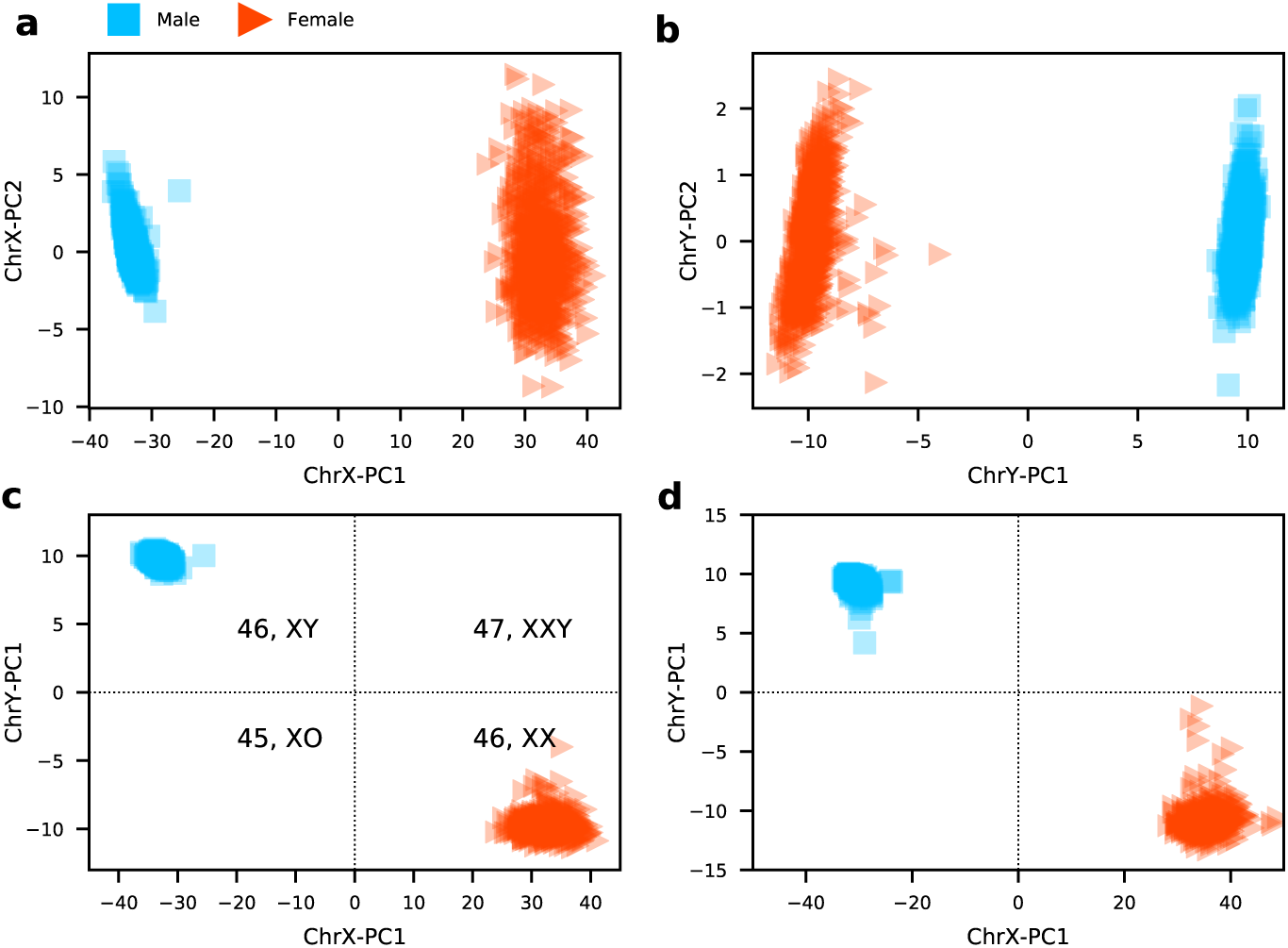
A sex classifier is constructed by applying two PCAs on two sex chromosomes separately. **a**: The first two components on ChrX. **b**: The first two components on ChrY. Results of **c** training set and **d** validation set produced by the sex classifier, all samples are classified into four categories: 46XY, 46XX, 47XXY, and 45XO.

Similarly, a PCA is performed using the 284 CpG sites of ChrY, and as that of ChrX, the first component accounted for the most variances can make a good separation between male and female samples (Fig. 2b). As the result of this, the first component can be used to divide samples into two categories: 1) with Y and 2) without Y.

Finally, the two first components of the two PCAs which both explained the most sex differences are utilized to build the sex classifier. Normal females have two X chromosomes and normal males have one X chromosome and one Y chromosome. By our sex classifer, male samples with 46,XY should locate in the top left area and female samples with 46,XX should distribute at the bottom right area (Fig. 2c). It is reasonable to suggest that this model can be applied to identify samples with sex aneuploidy: samples with 45,XO will be placed at the bottom left corner, and samples with 47,XXY should be distributed at the top right corner.

### Comparison with other tools

To compare the proposed sex classifier with three other existing sex prediction classifiers from DNA methylation microarray data taken from the R packages, *minfi* [13], *ewastools* [12] and *sEst* [14], we take GSE51032 [17] as a benchmark dataset, as it was used in developing *ewastools* and *sEst*. GSE51032 includes 857 samples (188 men and 657 women) and their source tissue are all from buffy coat. Fig. 4 shows the results generated by the four methods, as we can see, there are eight samples (four males and four females) displaying mismatches between predicted sex and labelled sex, and the mismatches are consistent in the results from four methods, thus we have high confidence that the eight samples are mislabelled. Two samples (marked by black circles) are identified by our classifier as 47,XXY, *sEst* also identified the two outliers. However, only one of the two samples appears as an outlier from *minfi* and *ewastools*, and the other one stays close with the male group.

**Figure 4.**
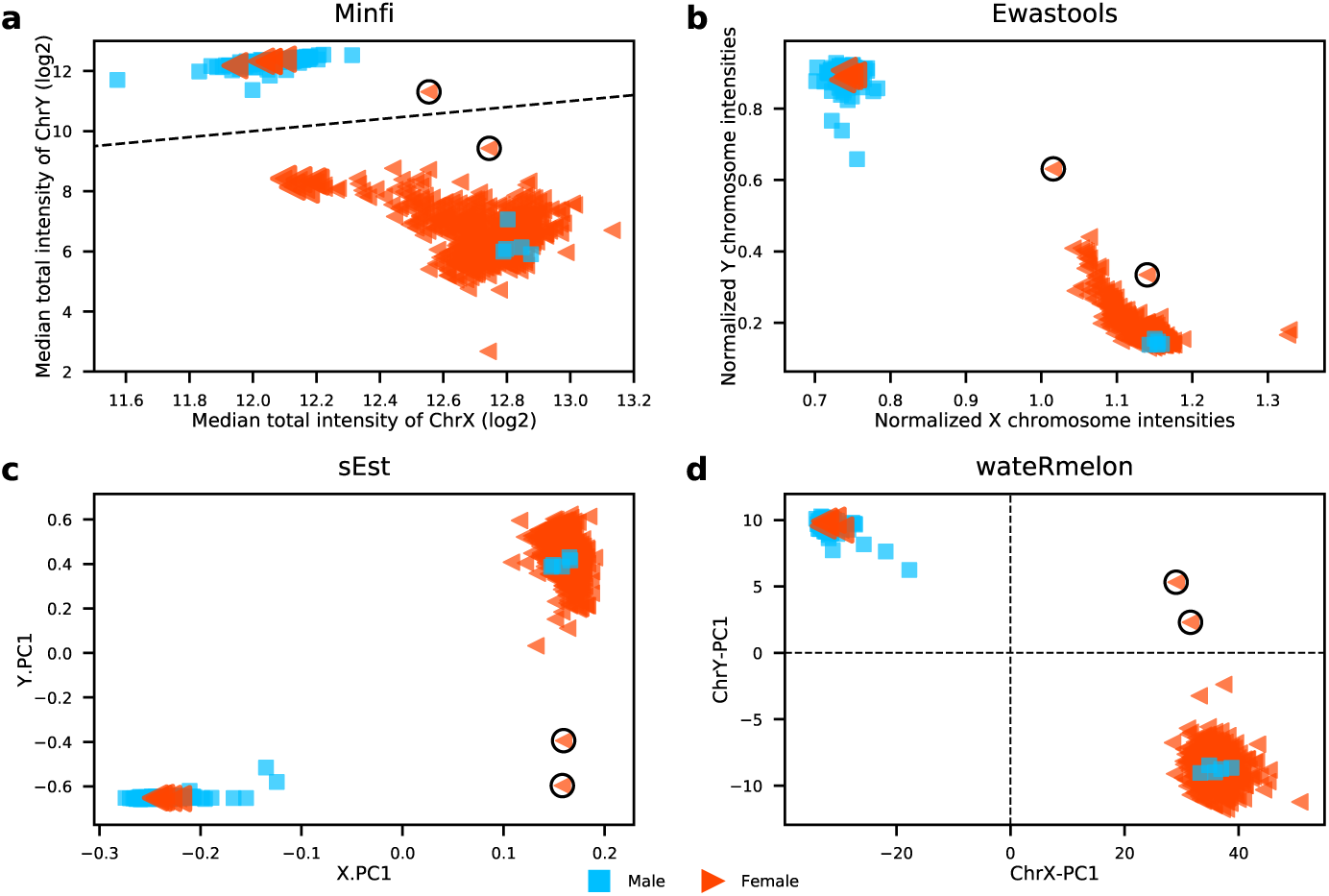
Comparisons of sex prediction ability between four tools. **a**. minfi, **b**. ewastools, **c**. sEst, **d**. our classifier in wateRmelon. Two outlier samples are marked by black circles, blue square represents male and red triangle denotes female.

In general, all four methods show good performance in clustering male samples, however the method from *minfi* performs much poorer in clustering female samples compare to the other three tools, as some females are not distinguishable from males in the x-axis. The female group of the result generate by *ewastools* exhibits long tail towards males; the sex prediction tools in *minfi* and *ewastools* are both based on signal intensity therefore they produce more similar results than the other two tools. Our sex classifier and the method from *sEst* are both beta value based, although the two methods utilised beta values very differently and *sEst* requires further *p*-value information, the patterns of their results are similar. Overall, compared to three other sex prediction tools, our proposed method is highly robust and shows better or similar performance in clustering females and males.

### Performance evaluation

The DNA methylation levels of samples from training set and validation set are assessed by 450k array and EPIC array respectively. As we can see from the results (Fig. 2), the proposed model has correctly classified all samples in the two datasets, proving that the proposed classifier is highly robust and compatible with both platforms.

The proposed sex classifier is trained and validated using whole blood samples. As whole blood is a heterogeneous collection of different cell types, to investigate whether our classifier is biased by blood cell types, we tested its performance on DNA methylation data derived from five purified blood cell types—B cells, CD4 T cells, CD8 T cells, monocytes and granulocytes from 28 individuals. As shown in Fig.3a and Fig.3b, all the five cell types are clustered into two sex groups and we could not find any or very minor differences between cell types. Collectively, these results suggest that the proposed sex classifier is robust to blood cell types.

**Figure 3.**
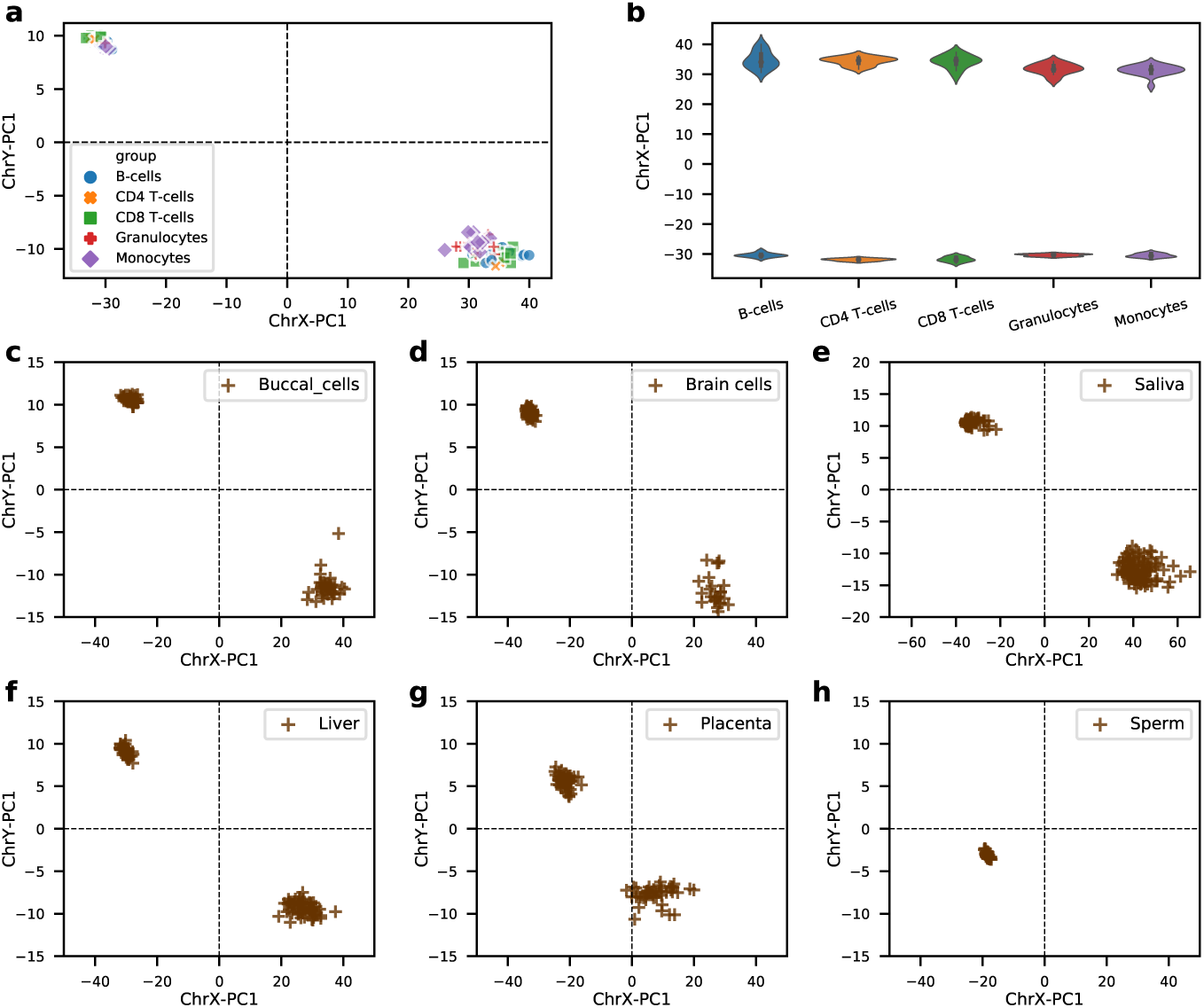
The sex classifier was evaluated across five blood cell types (a and b) and six other human tissues (c-h). **a**. Scatter plot showing results from five blood cell types: B cells, CD4 T cells, CD8 T cells, monocytes and granulocytes. **b**. On X chromosome, the five blood cell types showing similar results. **c**. Buccal cells; **d**. Brain cells; **e**. Saliva; **f**. Liver; **g**. Placenta; **h**. Sperms.

Although blood is the most studied tissue in EWAS, there are also many DNA methylation studies that use samples from other types of human tissue. To evaluate our sex classifier’s range of application, we further tested its performance on several other most studied human tissues, including saliva, buccal cells, brain cells, liver, placenta, and sperm. Results from Fig.3c to Fig.3f demonstrate that the proposed classifier is robust in these vastly different types of tissues—saliva, buccal cells, brain cells, and liver. However, despite we can observe two groups clustering within placenta, the females are much loosely clustered along the x-axis(Fig 3g). Additionally, boundaries have diverged greatly on both ChrX and ChrY, with this being more pronounced on ChrX.

Interestingly, sperm samples clustered into a single group, located in the bottom left region (Fig3.h). This area is typically recognised by our sex classifier as 45,XO. As sperm cells are a mixture of two types of haploid cells (23,X and 23,Y) this suggests that their methylation levels are lower on ChrY compared to other mature human tissues.

### Predicting sex chromosome aneuploidy

DNA methylation has been an important way to study the various developmental symptoms caused by copy number aberrations of the sex chromosome [18]. Earlier, we proposed that our classifier can be applied to identify samples with abnormal sex chromosomes, including 45,XO and 47,XXY. To further validate its ability, we searched the public repositories for positive samples with clinical diagnosis. As a result of this, we obtained five cases (Table 1) diagnosed as Turner syndrome from two studies [19, 20]. As hoped, they are all clearly classified as 45,XO by our model (Fig. 5), proving our classifier’s ability to predict females with only one X chromosome.

**Table 1.**
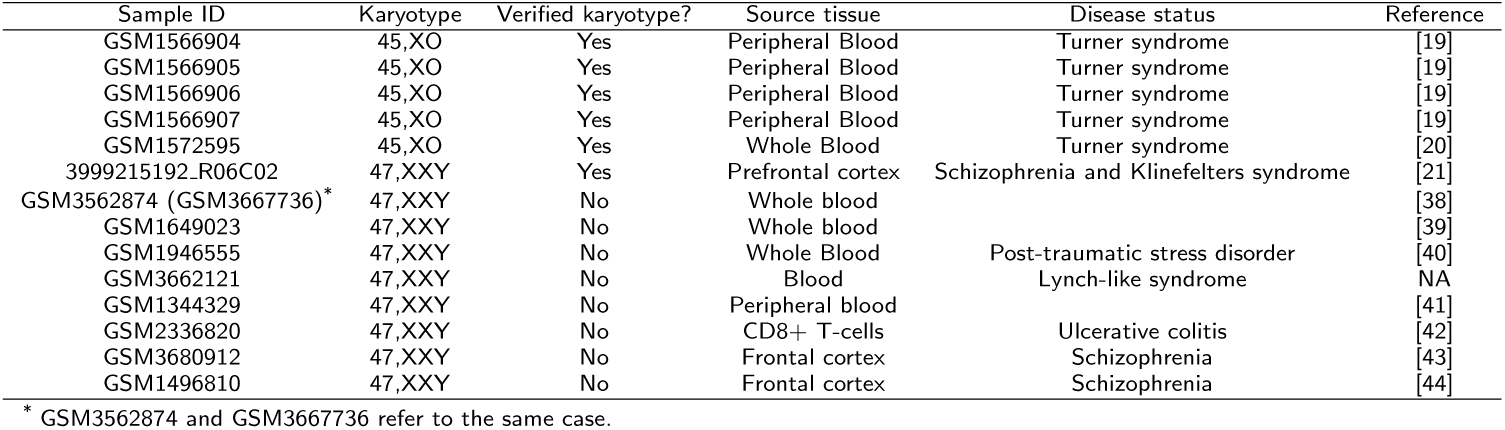
Samples with verified or suspect abnormal karyotypes from GEO.

**Figure 5.**
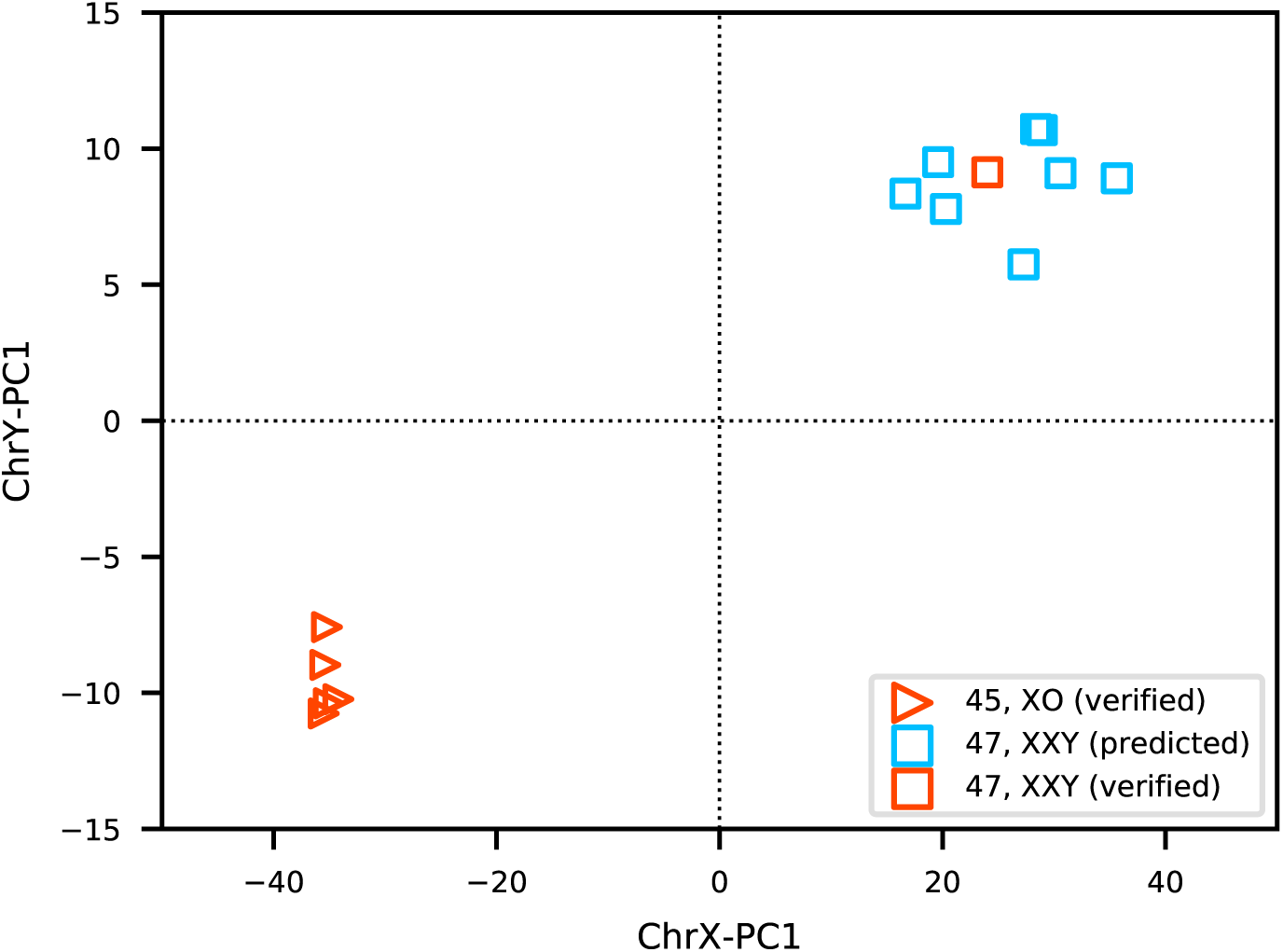
The proposed classifier is verified its ability to predict sex chromosome aneuploidy in five Turner syndrome samples and one Klinefelter syndrome case, it also predicted eight potential 47,XXY cases from GEO.

Viana et al. reported a male with schizophrenia carrying an extra X chromosome [21] which is also clearly classified as 47,XXY by our method (Fig. 5). Unfortunately, we did not find any publicly available samples from those diagnosed with Klinefelter syndrome. Unlike Turner syndrome, most patients with Klinefelter syndrome have only mild symptoms and are never diagnosed. It is interesting to find out how many samples in GEO having a karyotype of 47,XXY but not linked to a diagnosis. By applying our classifier to scan the GEO datasets, we find a total of eight samples (Table 1) which are highly likely to be 47,XXY (Fig. 5). It should be noted that we only include these samples sourced from blood or brain cells related tissues and their DNA methylation level are assessed by 450K or EPIC arrays; we also do not include those samples which located near the boundaries which may be low-level sex chromosome mosaics (46,XX/47,XXY). It is interesting that two of the eight suspect abnormal samples are from those diagnosed with schizophrenia. Martin et al. found that Klinefelter patients have nearly a four times higher risk of schizophrenia [22], which accords with our observation of more 47,XXYs with schizophrenia. Studying the methylation patterns of these syndromes will provide more insights into these diseases.

## Discussion

There are two principal reasons to require a good and simple sex classifier based on methylation data. First, there are still many samples in GEO that do not have sex annotations, thus an accurate classifier can provide reliable sex information. Second, due to data entry errors, there are non-negligible proportions of mislabelled samples in the public database. A mismatch between reported sex and predicted sex would be a clear indication of a wrong annotation and introduces doubt on the accuracy of the rest of the phenotype information for that sample, hence it is reasonable to remove these mislabelled samples before downstream analyses. We would recommend sex checking to be a standard part of all DNA methylation QC pipelines. Here in this study, the proposed sex classifier is straightforward and the outcomes are highly intuitive.

In this study, we first obtained a group of significant sex-associated CpG sites. 90% of these located on the X chromosome are more methylated in females than that in males, this is mainly due to the effect of X-chromosome inactivation: one of the two X chromosomes in females is randomly chosen for inactivation (highly methylated) to balance the extra gene expression dosage [23, 24]. This also justified that our classifier was built on blood samples could work well across a wide range of other tissue types.

The proposed sex classifier shows robust performance across a wide range of tissue types despite it is built upon whole blood samples. We choose blood samples because they are easily accessible and are the most widely used tissue for measuring DNA methylation and have been adopted in most large cohort studies. However, whole blood is a heterogeneous collection of different cells, and their cell composition changes across age [25]. Different cell types can have distinct methylation profiles even though they share identical genetic makeup [26]. Here as our results have shown that the proposed model is not biased among different blood cell types; we also demonstrated the proposed classifier performs well across a wide range of human tissues, including saliva, buccal cells, brain cells, liver. These results suggest that our model is not driven by blood-specific sex differences, but it has captured the more general sex-associated differences across human tissues and cell types. However, we have also found some tissues such as placenta (Fig3.h) showing an ambiguous boundary between the two sexes. Placenta is a fetal-maternal endocrine organ responsible for ensuring proper fetal development throughout pregnancy [27]. The fetal part of the placenta has the same genetic composition as fetus, whereas it exhibits apparent different DNA methylation patterns. Our results demonstrate placenta samples are less distinguishable between the two sex groups, showing both ChrX in female placentas and ChrY in male placentas are less methylated than that in other normal tissues. During the early development of human embryo, sperm cells are highly methylated and then become hypomethylated after fertilization [28]. Our results have shown that those sex-associated CpG on X chromosomes of sperm cells exhibited similar methylation patterns with other normal male tissues, however, the Y chromosomes are much less methylated. Collectively, our method can also be used to evaluate the methylation level of the two sex chromosomes in different tissues.

Our method can be readily applied to almost all DNA methylation datasets in GEO. A large portion of DNA methylation datasets uploaded to GEO are not in IDAT format, which is prerequisite by using *minfi* and *ewastools*, many of these datasets only include density values of the methylated and unmethylated signals. Our sex classifier developed in this paper is based on beta values of those differently methylated CpG loci between the two sexes, users are only required to feed the whole beta value matrix, which can be easily computed from the density files, to the ‘estimateSex’ function in *wateRmelon* to obtain final sex predictions.

The underlying mechanism of our sex classifier is very intuitive: females have higher levels of methylation on ChrX, on the contrary, males are less methylated on ChrX and show strong methylation signals on ChrY. We have also demonstrated that the proposed classifier can be applied on both 450K and EPIC arrays. Compared to signal density-based methods such as *minfi* and *ewastools*, the methylation ratio-based method from our sex classifier and *sEst* provide better separation between the two sexes (Fig 4). In addition, both *minfi* and *ewastools* require at least one female and one male in the input samples to make correct sex predictions, however, our method and *sEst* do not have a such limitation. In term of running speed, our method is more than four times faster than *sEst* when the number of input samples exceeds 1,000.

We have provided a powerful tool that can identify sex chromosome aneuploidies (45,XO and 47,XXY) from DNA methylation data. This function has been verified in five Turner syndrome samples and one Klinefelter syndrome case, we should acknowledge that we need much more positive cases to testify its sensitivity and specificity. It is a pity that we did not find any DNA methylation samples labelled as Klinefelter syndrome in the public repositories. Nevertheless, we found eight cases in the GEO database with great potential to be 47,XXY by applying our classifier, with the knowledge that most patients with Klinefelter syndrome have only mild symptoms and are never diagnosed. Those eight suspect Klinefelter syndrome cases can be good candidates to study the various developmental symptoms caused by copy number aberrations of sex chromosomes.

## Conclusion

In this study, we constructed a very intuitive sex classifier, simply based on the most robust CpG sites on the sex chromosomes, which not only can be used for sex predictions but also applied to identify samples with sex chromosome aneuploidy. Our classifier has been integrated into the *wateRmelon* Bioconductor package, which is freely and easily accessible by calling the ‘estimateSex’ function.

## Methods

All statistical analyses were conducted either by R (version 3.6.0, https://www.r-project.org/) or Python (version 3.7.4, https://www.python.org/). Raw data for all datasets were processed by bigmelon package [29], methylated and unmethylated signals were extracted from idat files and beta values are calculated as:

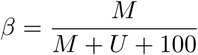

where *β* is beta value, *M* denotes methylated densities and *U* represents unmethylated densities.

## Availability of data

All the DNA methylation data except for the validation set are public available and were obtained from the GEO public repository. The training set is from GSE105018[30] which includes 832 male and 826 female whole blood samples, the validation set which includes 1175 whole blood samples is available from the European Genome-phenome Archive under accession EGAS00001002836 (https://www.ebi.ac.uk/ega/home). Other datasets: purified blood cell types (GSE103541 [31]), buccal cells (GSE137884 [32]), brain cells (GSE112179 [33]), saliva (GSE78874 [34]), liver (GSE119100 [35]), placenta (GSE100197 [36]), sperms (GSE64096 [37]). The one Klinefelter syndrome positive sample is available upon request. More details about these datasets are shown in Table 2.

**Table 2.** Summary of datasets used in this study.

| Dataset | Source | Platform | Number | Male/Female | Age(years) | Reference |
| --- | --- | --- | --- | --- | --- | --- |
| GSE105018 | Whole blood | 450k | 1658 | 832/826 | 18 - 18 | [30] |
| UKHLS | Whole blood | EPIC | 1175 | 489/686 | 28 - 98 | [45] |
| GSE103541 | Purified blood cells | EPIC | 145 | NA | NA | [31] |
| GSE137884 | Buccal cells | 450k | 89 | 51/38 | 3 - 6 | [32] |
| GSE112179 | Brain cells | EPIC | 100 | 75/25 | 23 - 77 | [33] |
| GSE78874 | Saliva | 450k | 259 | 146/113 | 36 - 88 | [34] |
| GSE119100 | Liver | EPIC | 108 | 46/62 | 25 - 71 | [35] |
| GSE100197 | Placenta | 450k | 102 | NA | NA | [36] |
| GSE64096 | Sperms | 450k | 40 | NA | NA | [37] |
| GSE51032 | Buffy coat | 450k | 845 | 188/657 | 34 - 72 | [17] |

## Author’s contributions

YW designed the method, wrote the codes and performed all the analysis. YW wrote the paper with contributions from LCS, XZ and KM. EH, OAG, TJG, MK and JM provided critical insights. XZ, KM and LCS advised and oversaw the work. All authors read and approved the final manuscript.

## Acknowledgements

The authors acknowledge the use of the High Performance Computing Facility (Genome) and its associated support services at the University of Essex in the completion of this work.

